# Influenza A virus environmental persistence is driven by the hemagglutinin and the neuraminidase

**DOI:** 10.1101/268722

**Authors:** Thomas Labadie, Christophe Batéjat, Jean-Claude Manuguerra, India Leclercq

## Abstract

The transmission routes of Influenza A viruses (IAVs) submit virus particles to a wide range of environmental conditions that affect their transmission. In water, temperature, salinity and pH are important factors modulating viral persistence in a strain-dependant manner, and the viral factors driving IAV persistence remained to be described. We used an innovative method based on a real-time cell system analysis to quantify viral decay in an environmental model. Thus, we identified the viral hemagglutinin (HA) and neuraminidase (NA) as the main proteins driving the environmental persistence by comparing the inactivation slopes of several reassortant viruses. We also introduced synonymous and non-synonymous mutations in the HA or in the NA that modulated IAV persistence. Our results demonstrate that HA stability and expression level, as well as calcium-binding sites of the NA protein are molecular determinants of viral persistence.

Finally, IAV particles could not trigger membrane fusion after environmental exposure, stressing the importance of the HA and the NA for environmental persistence.

## Introduction

Influenza A viruses (IAVs) has a wide host range, which allows them to spread almost everywhere on the planet. In nature, the H1N1 subtype infects several hosts such as domestic and aquatic birds, humans, swine or dogs and thus spreads through a wide range of environmental conditions. In humans, a pandemic H1N1 virus emerged in 2009 after the reassortment between a swine H1N1 virus from the Eurasian lineage and a swine H2N1 virus descended from a triple reassortment between a swine H1N1 virus from the North American lineage, an avian virus and the human H3N2 virus. In aquatic birds, which are the main reservoir of these viruses, IAVs spread mainly by a fecal-oral route through water. In poultry such as chicken and turkeys, or in mammalian species, IAVs mainly have a respiratory tropism, and the virus spreads by contact between infected and susceptible hosts or by contaminated fomites, as well as through aerosol or respiratory droplets. In any case, the transmission routes of IAVs submit virus particles to a wide range of environmental conditions, which more or less rapidly affect them. In water, the time for IAV inactivation depends on widely studied abiotic factors such as temperature (Brown et al. 2009, 2007; Dublineau et al. 2011; Keeler et al. 2014; Nazir et al. 2010; Poulson et al. 2016; D.E Stallknecht and Brown 2009; H. Zhang et al. 2014), water salinity (Brown et al. 2009; Dublineau et al. 2011; Keeler et al. 2014; Poulson et al. 2016) and pH (Brown et al. 2009; Keeler et al. 2014; Poulson et al. 2016) and can range from a few days in saline water (35 g.L^−1^ NaCl) at 35°C to several years at 4°C (Dublineau et al. 2011). The environment acts both as a reservoir (Roche et al. 2014; David E. Stallknecht et al. 2010) and as a bottleneck for IAVs evolution. It was experimentally demonstrated that IAV persistence in water is extremely variable among strains and subtypes (Dublineau et al. 2011; Lebarbenchon et al. 2012; D.E Stallknecht and Brown 2009). However, inactivation processes of avian IAVs in the environment are far from being well understood (Lebarbenchon et al. 2010). Only a few viral factors, such as the viral envelope origin and composition (Shigematsu et al. 2014; Bajimaya et al. 2017) and the pH of activation of the HA (Reed et al. 2010) are known to affect virus survival. On the other hand, the viral envelope and the viral genome are affected much more slowly than the actual loss of infectivity in water, suggesting that other viral factors drive IAV survival (Shigematsu et al. 2014; Dublineau et al. 2011). The envelope of IAV structure is made of two external glycoproteins, the hemagglutinin (HA) and the neuraminidase (NA), and a proton channel M2, which are inserted into a lipid bilayer surrounding a matrix M1 protein layer. Inside viral particles, the segmented and negative strand RNA genome surrounded by a nucleocapsid (NP) interacts with the viral polymerase made of the three proteins PB1, PB2 and PA to constitute the ribonucleoprotein (RNP) complex. The 8 RNPs interact closely with the M1 proteins.

In order to identify viral genetic drivers of the persistence phenotype, we generated reassortants from H1N1 viruses, which do not have the same persistence in an environmental model, and compared their inactivation slopes. As a model, we used saline (35 g.L^−1^ NaCl) water at 35°C, because it is the average salt concentration in the ocean and this temperature allows observing differences of viral persistence more rapidly. Using fluorescence imaging microscopy, we also wanted to understand how our environmental model disrupts viral functions and infectivity of IAVs. We identified molecular determinants of the persistence phenotype in the environment, by introducing codon-optimized synonymous mutations or non-synonymous mutations in the HA or the NA gene. Altogether, our results demonstrate that the survival of Influenza viral particles is predominantly driven by the two external glycoproteins, and that environmental conditions affect the HA mediated steps during viral entry.

## Results

### Influence of viral proteins on H1N1 strains persistence outside the host

Influenza A viruses, such as the seasonal H1N1 virus of 1999 and the pandemic H1N1 virus of 2009, show different persistence in water (Dublineau et al. 2011; Lebarbenchon et al. 2012; D.E Stallknecht and Brown 2009). This study aimed at characterizing viral molecular drivers associated with variations of persistence in saline water (35 g.L^−1^ NaCl) at 35°C used as an environmental model. These temperature and salinity conditions were selected because they allow observing variations between strains in a short period of time (Dublineau et al. 2011), and 35 g.L^−1^ NaCl is the average salt concentration in the oceans (Millero et al. 2008). In order to understand how viral proteins influence the persistence of H1N1 influenza particles, reassortant viruses between a pandemic H1N1 virus, the A/Bretagne/7608/2009 strain (whole/2009) and a pre-pandemic H1N1 virus, the A/WSN/1933 strain (whole/1933) were generated using a plasmid-based reverse genetics system. We rescued reassortant viruses containing either the HA/NA segments, the PB1/PB2/PA segments, the M segment or the NS/NP segments from A/Bretagne/7608/2009 strain with the other segments from A/WSN/1933 as a complementary genomic backbone. We conversely rescued the same mirroring set of viruses with the same pool of segments but from the A/WSN/1933 strain and with the genomic backbone constituted by A/Bretagne/7608/2009 virus segments. The composition of each virus is detailed in tables 1 and 2. All reassortants grew except the NS-NP/1933 virus. Thus, we generated seven different reassortant viruses and compared their persistence with a real-time cell analysis system (RTCA), as described in the Methods section (Fig. 1A). To quantify viral decay, we monitored CIT_50_ values after 1, 24 and 48 hours of exposure in saline water at 35°C. The increase of CIT50 value over time reflects the loss of infectivity and the progressive inactivation of viral particles (Fig. 1B). We calculated inactivation slopes from experimental CIT_50_ values (Fig. 1C). The mean inactivation slope of the whole/2009 virus in saline water at 35°C was 4.4 CIT_50_.day^−1^ (Fig. 1D), which is twice more stable than the whole/1933 virus that has a mean inactivation slope of 8.3 CIT_50_.day^−1^ (Fig. 1E). When compared with the whole/2009 virus (Fig. 1D), replacing the 2009 HA and NA by the 1933 HA and NA (HA-NA/1933 virus), or the 2009 M segment by a 1933 M segment (M/1933 virus) significantly destabilized the virus with mean inactivation slopes of 6.9 and 11.3 CIT_50_.day^−1^ respectively. On the contrary, the Pol/1933 virus persistence was not significantly different from that of the whole/2009 virus, with a mean inactivation slope of 3.3 CIT_50_.day^−1^. Compared with the whole/1933 virus (Fig. 1E), replacement of the 1933 HA and NA by the 2009 HA and NA (HA-NA/2009 virus) significantly increased the persistence of the virus with a mean inactivation slope of 6.7 CIT_50_.day^−1^. The replacement of the 1933 M or NS and NP segments by their 2009 counterparts (M/2009 and NS-NP/2009 viruses) did not change significantly their persistence. More surprisingly, replacing the polymerase PB1, PB2 and PA segments of 1933 by the polymerase segments of 2009 (Pol/2009 virus) significantly destabilized this reassortant virus with a mean inactivation slope of 12.4 CIT_50_.day^−1^.

**Table 1.** Engineered reassortants between A/Bretagne/7608/2009 H1N1 virus and A/WSN/1933 H1N1 virus.

**Tables and figures legends**
| Reassortant name | A/Bretagne/7608/2009 | A/WSN/1933 |
| --- | --- | --- |
| Whole/2009 | HA, NA, M, PB1, PB2, PA, NS, NP | - |
| HA-NA/2009 | HA, NA | M, PB1, PB2, PA, NS, NP |
| M/2009 | M | HA, NA, PB1, PB2, PA, NS, NP |
| Pol/2009 | PB1, PB2, PA | HA, NA, M, NS, NP |
| NS-NP/2009 | NS, NP | HA, NA, M, PB1, PB2, PA |
| Whole/1933 | - | HA, NA, M, PB1, PB2, PA, NS, NP |
| HA-NA/1933 | M, PB1, PB2, PA, NS, NP | HA, NA |
| M/1933 | HA, NA, PB1, PB2, PA, NS, NP | M |
| Pol/1933 | HA, NA, M, NS, NP | PB1, PB2, PA |

**FIG 1.**
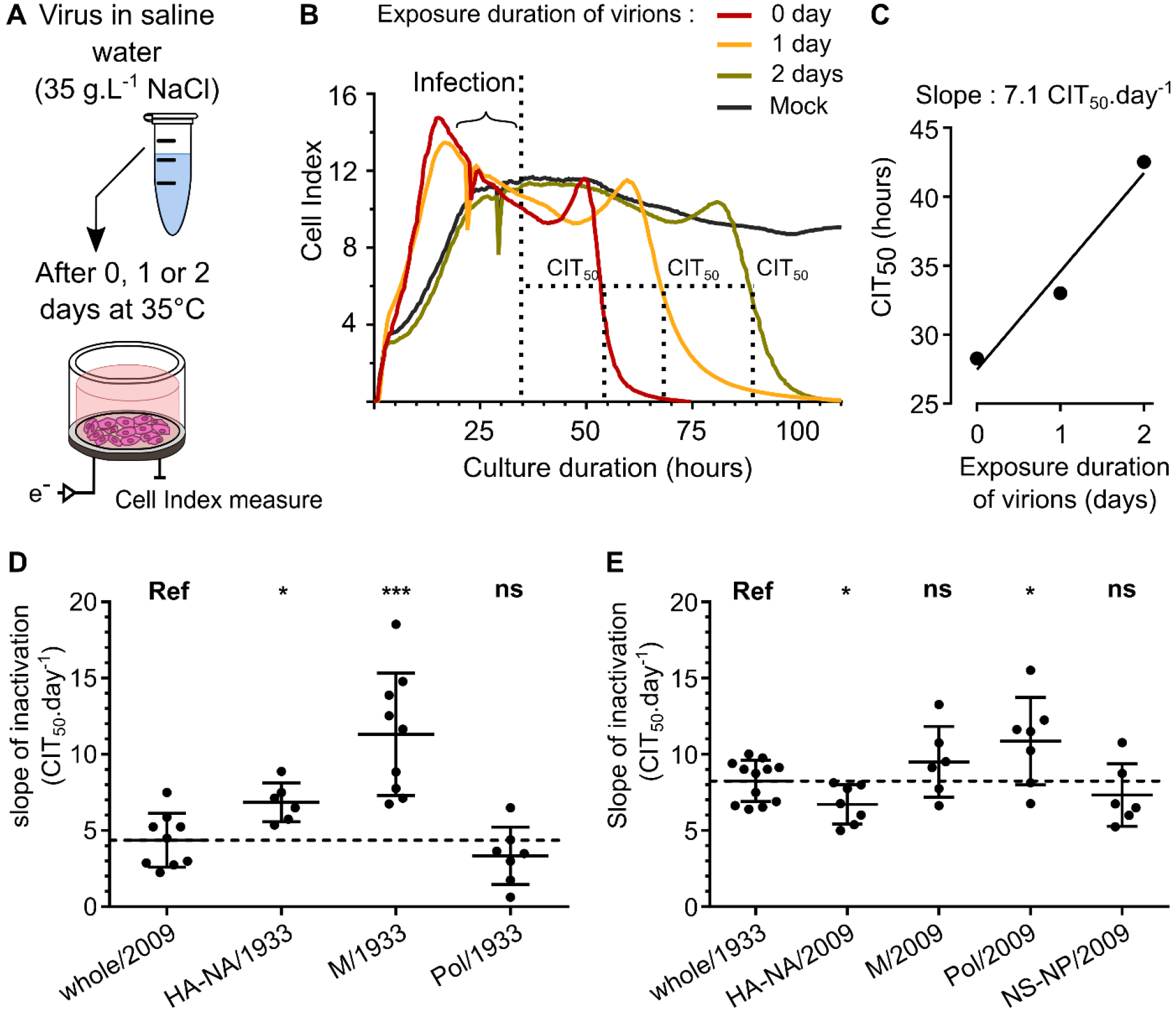
Influence of viral proteins on H1N1 viruses persistence in saline water at 35°C. (A) Viral particles were diluted in saline water (35 g.L^−1^ NaCl) and exposed to a temperature of 35°C for 0, 1 or 2 days. MDCK cells were infected by exposed viruses and impedance was monitored continuously and plotted as “Cell Index” (CI). (B) CI decrease due to virus-induced cytopathogenic effect was quantified with the CIT_50_ value, which is the necessary time in hours to measure a 50% decrease from the initial CI, always set as the cell index value 5 hours after infection (symbolized as a vertical dashed line). (C) The CIT_50_ values of an exposed virus increase with the duration of exposure in saline water at 35°C. It reflects the loss of infectivity and a linear regression analysis allowed calculatingthe viral inactivation slope. (D) and (E) Inactivation slopes comparison between (D) the 2009 wild-type H1N1 virus (reference; whole/2009) and reassortant viruses harbouring different genomic segments of the 1933 H1N1 strain or (E) between the 1933 wild-type H1N1 virus (reference; whole/1933) and the reassortant viruses harbouring genomic segments of the 2009 H1N1 strain. Means of inactivation slopes (horizontal lines) were compared using a Wilcoxon-Mann-Whitney test (NS, *P*>0.1; \**P*<0.05; ***P<0.0005). Each dot represents a single inactivation slope (d and e) and vertical lines represent the standard deviation. Dashed lines represent the mean inactivation slope of the virus used as a reference for comparison.

Thus, once the HA and NA were isolated from their genomic context into a new virus, the persistence phenotype of this new virus tended to be the same as the virus from which HA and NA were originated. The matrix protein and to a certain extent the viral polymerase only induced a decrease of viral persistence without reflecting the phenotype of their original virus. Altogether, these results suggested that HA and NA are the most important viral proteins driving the persistence phenotype.

### Molecular determinants of the HA driving environmental persistence

In order to understand how the HA protein modulates environmental persistence, we rescued two reassortant viruses bearing a HA and NA either from the H1N1 A/NewCaledonia/20/1999 strain (HA-NA/1999) or from the pandemic A/Paris/2590/2009 strain (HA-NA/2009) (Table 2), both with the same genomic backbone belonging to the A/WSN/1933 strain. We selected these HA and NA because the two strains have a different persistence phenotype in saline water at 35°C (Dublineau et al. 2011) in favour of the pandemic strain. Using a third strain also allowed us to compare the HA and NA of the 2009 pandemic strain and the pre-pandemic strain independently of their parental genomic context. We wanted to assess whether introducing changes in the HA sequence had an impact on viral persistence. We thus synthesized a codon-optimized gene encoding the 1999 HA (HA_opti_), producing the same HA but containing 24.4% of synonymous substitutions. We then generated the corresponding reassortant virus bearing this HAopti and the 1999 NA in the A/WSN/1933 backbone (HA_opti_-NA/2009, Table 2). The slope of inactivation was significantly lower in saline water at 35°C for the HA-NA/2009 virus compared with the HA-NA/1999 virus inactivation slope (Fig. 2A), respectively 5.9 and 9.9 CIT_50_.day^−1^. This result is in agreement with that of a previous study (Dublineau et al. 2011) on the persistence phenotype of the wild-type non-reassortant A/Paris/2590/2009 and the A/NewCaledonia/20/1999 strains, and confirms that HA and NA are indeed the main viral factors leading the persistence of viral particles. Interestingly, HA_opti_-NA/1999 was as stable as HA-NA/2009 and thus significantly more stable than HA-NA/1999 (Fig. 2A) despite the identical amino acid composition of HA.

**Table 2.** Engineered reassortants between A/Paris/2 590/2009 H1N1 virus and A/NewCaledonia/20/1999 H1N1 virus with the A/WSN/1933 H1N1 virus used as a genomic backbone

| Reassortant name | A/Paris/2590/2009 | A/NewCaledonia/20/1999 | A/WSN/1933 |
| --- | --- | --- | --- |
| HA-NA/2009 | HA, NA | - | M, PB1, PB2,<br>PA, NS, NP |
| HA-NA/1999 | - | HA, NA |  |
| HA<sub>opti</sub>-NA/1999 | - | HA (codon optimized), NA |  |

**FIG 2.**
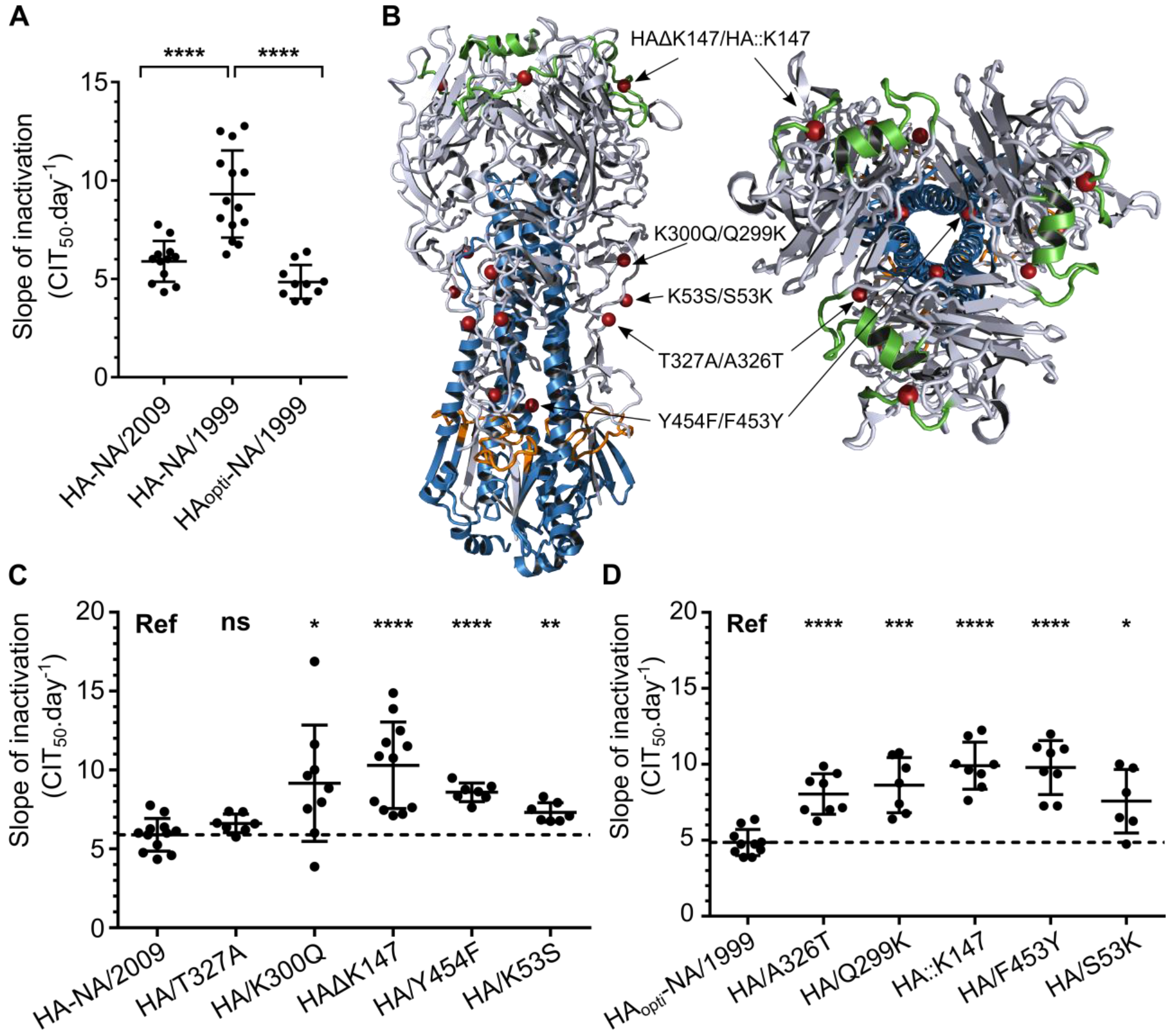
Synonymous and non-synonymous substitutions in the HA change environmental persistence. (A) Inactivation slopes of reassortant viruses bearing a HA and a NA either from 2009 (HA-NA/2009), 1999 (HA-NA/1999) or 1999 with 24% of the HA mutated by synonymous substitutions (HA_opti_-NA/1999). Each dot represents a single inactivation slope and their means (horizontal lines) were compared using an ANOVA test (NS, *P*>0.05; \*\*\*\**P*<0.0001). **(B)** Selected mutations either on the HA of 2009 or the optimized HA of 1999 are indicated by red spheres and arrows on the crystal structure of a 2009 H1N1 virus trimeric HA (3LZG) (side or top-down views). (C) and (D) Mean inactivation slopes of mutated viruses were compared to the HA-NA/2009 virus (C) or HA_opti_-NA/1999 virus (D). Each dot represents a single inactivation slope value and their means (horizontal lines) were compared using a Wilcoxon-Mann-Whitney test (NS, *P*~0.05; \**P*<0.02; \*\**P*<0.005; \*\*\**P*=0.0001; \*\*\*\**P*<0.0001). Vertical lines represent the standard deviation and dashed lines represent the mean inactivation slope of the virus used as a reference for comparison.

We also introduced non-synonymous mutations in the HA nucleotide sequence to study the impact of a single HA amino-acid change on virus persistence. For this purpose, we compared the HA amino-acid sequences of the HA-NA/1999 and HA-NA/2009 viruses and, based on a preliminary mutation screening performed using a lentiviral pseudo-particles system (Sawoo et al. 2014), we selected five residues in the HA protein that might influence virus particle persistence outside the host (Fig. 2B). Ten reassortant viruses bearing HA with amino-acid substitutions or insertions/deletions were generated after site-directed mutagenesis of the pPol-HA plasmid. We then exposed these mutated HA-NA/2009 and HA_opti_-NA/1999 viruses to saline water at 35°C (Fig. 2C and 2D). The HA/Y454F substitutions or HAΔK147 deletion in the HA of HA-NA/2009 virus induced a significant increase of the mean inactivation slope, respectively to 9.2 and 10.3 CIT_50_.day^−1^, thus generating very unstable mutants. Substitutions HA/K300Q and HA/K53S also destabilized the HA-NA/2009 virus but to a lesser extent, whereas the HA/T327A substitution did not affect its persistence, with mean inactivation slopes of 9.2, 7.3 and 6.6 CIT_50_.day^−1^ respectively. The insertion of HA::K147 or substitutions A326T, F453Y and Q299K in the HA of HA_opti_-NA/1999 virus greatly decreased the persistence of the virus, with a mean inactivation slope of 9.9, 8.0, 9.8 and 9.6 CIT_50_.day^−1^, respectively. The S53K substitution however had less impact on the persistence, with a mean inactivation slope of 7.6 CIT_50_.day^−1^. Altogether, these results demonstrate that synonymous mutations or a single amino acid change in the HA is sufficient to negatively affect the viral persistence outside the host.

### Viral particles cannot trigger membrane fusion after being exposed to saline water at 35°C

Since the loss of hemagglutination titre of viral particles is slower than their loss of infectivity (Chu 1948) (Fig. 3A and Fig. S1), we decided to evaluate whether viruses were still able to bind their cellular receptor. For this purpose, we immuno-labelled the viral nucleoprotein (NP), which encapsidates the viral genome to form the ribonucleoprotein, and used confocal microscopy for its detection in infected MDCK cells. In cells infected with non-exposed HA-NA/2009 or HA-NA/1999 viruses, the NP protein was concentrated within the nucleus 2 hours after infection (Fig. 3B). On the contrary, in MDCK cells infected with exposed HA-NA/2009 or HA-NA/1999 viruses for 5 days to saline water at 35°C, we detected the NP protein close to the cell membrane but not in the nucleus after 2 hours of infection. We confirmed that NP localization of exposed viruses was similar to that of non-exposed viruses after 20 min of infection, when virus entry is not completely achieved (Fig. S2). We thus assessed the HA-triggered fusion of the viral membrane in infected MDCK cells using R18-labelled viruses at self-quenched concentration, a widely used technique to detect fusion of the viral membrane with the endosomal membrane (Pohl, Edinger, and Stertz 2014; Schelker et al. 2016). Fusion events were detected 20 min after infection by confocal microscopy in cells infected with viruses exposed 5 days to saline water at 35°C or non-exposed viruses (Fig. 3C). We detected almost no R18 dequenching signal in cells infected with HA-NA/2009 or HA-NA/1999 exposed viruses, whereas we observed numerous fusion events in cells infected with the non-exposed viruses. This result demonstrates that the loss of infectivity of H1N1 viruses in saline water at 35°C is a consequence of the HA inability to trigger membrane fusion.

**FIG 3.**
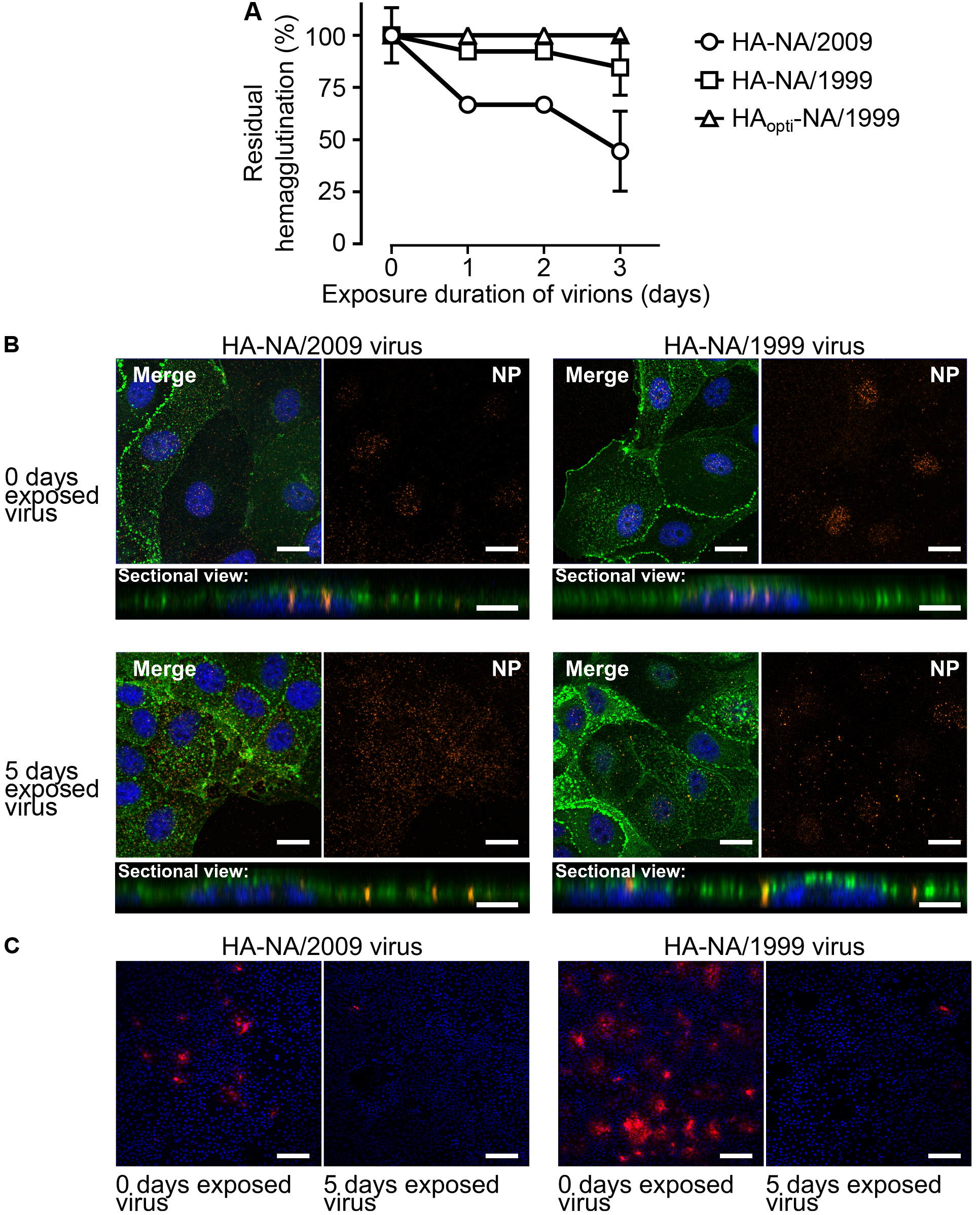
The HA of virus particles is still able to bind its cellular receptor but cannot trigger membrane fusion after being exposed. (A) Evolution of the residual hemagglutination titre (%) during exposure of reassortant viruses in saline water at 35°C. Vertical lines represent the standard deviation (N=3). (B) Confocal immunofluorescence microscopy (x40) of (A) MDCK cells 2 hours after infection by either the HA-NA/2009 virus (left), or the HA-NA/1999 virus (right), which have been exposed (bottom) or not (top) for 5 days to saline water (35 g.L^−1^ NaCl) at 35°C. Signal of the influenza NP immunolabelling is shown in orange. WGA labelling signal at the cell membrane is shown in green and the nucleus labelling with Hoescht dye is shown in blue. For each condition a merge of all channels (top left), the NP immunolabelling acquisition channel only (top right) or a Z-stack of sectional views at a distance of 0.42 μM (bottom) are shown. (C) MDCK cells (x10 magnification) 20 min after infection with both HA-NA/2009 and HA-NA/1999 viruses previously labelled with the R18 fluorophore at a self-quenched concentration. R18 dequenching signal due to the fusion of the viral envelope with the endosomal membrane is shown in red. Scale bars represent 20 μm (B, merged and NP views), 10 μm (B, sectional views) and 100 μm (C).

### HA stability at low pH is not the only determinant of viral persistence in water

A link between HA stability at low pH and stability to increasing temperatures has already been proposed (Imai et al. 2012; Krenn et al. 2011). Therefore, we measured the persistence of HA-NA/2009, HA-NA/1999 and mutated viruses in various pH-adjusted PBS solutions for 1 hour at room temperature (Fig. 4A). In addition, we quantified HA cell surface expression by flow cytometry in MDCK cells at 24 hours post infection (Fig. 4B). The most unstable viruses (HA/Y454F and HAΔK147 viruses; Fig. 2C) showed a lower HA surface expression level (Fig. 4B) and had a higher sensitivity at low pH compared with HA-NA/2009 virus. For the stable HA_opti_-NA/1999 virus, we observed a higher HA surface expression level compared with the unstable HA-NA/1999 virus, but they both had a high sensitivity at low pH. Similarly, the whole/1933 virus, which was more stable than the Pol/2009 virus (Fig. 1E) even if they both have the same HA and NA, had much higher HA expression levels in infected cells (Fig. 4C). Altogether, our results support the idea that HA expression level is a viral driver of environmental persistence when the HA has a high pH of inactivation.

**FIG 4.**
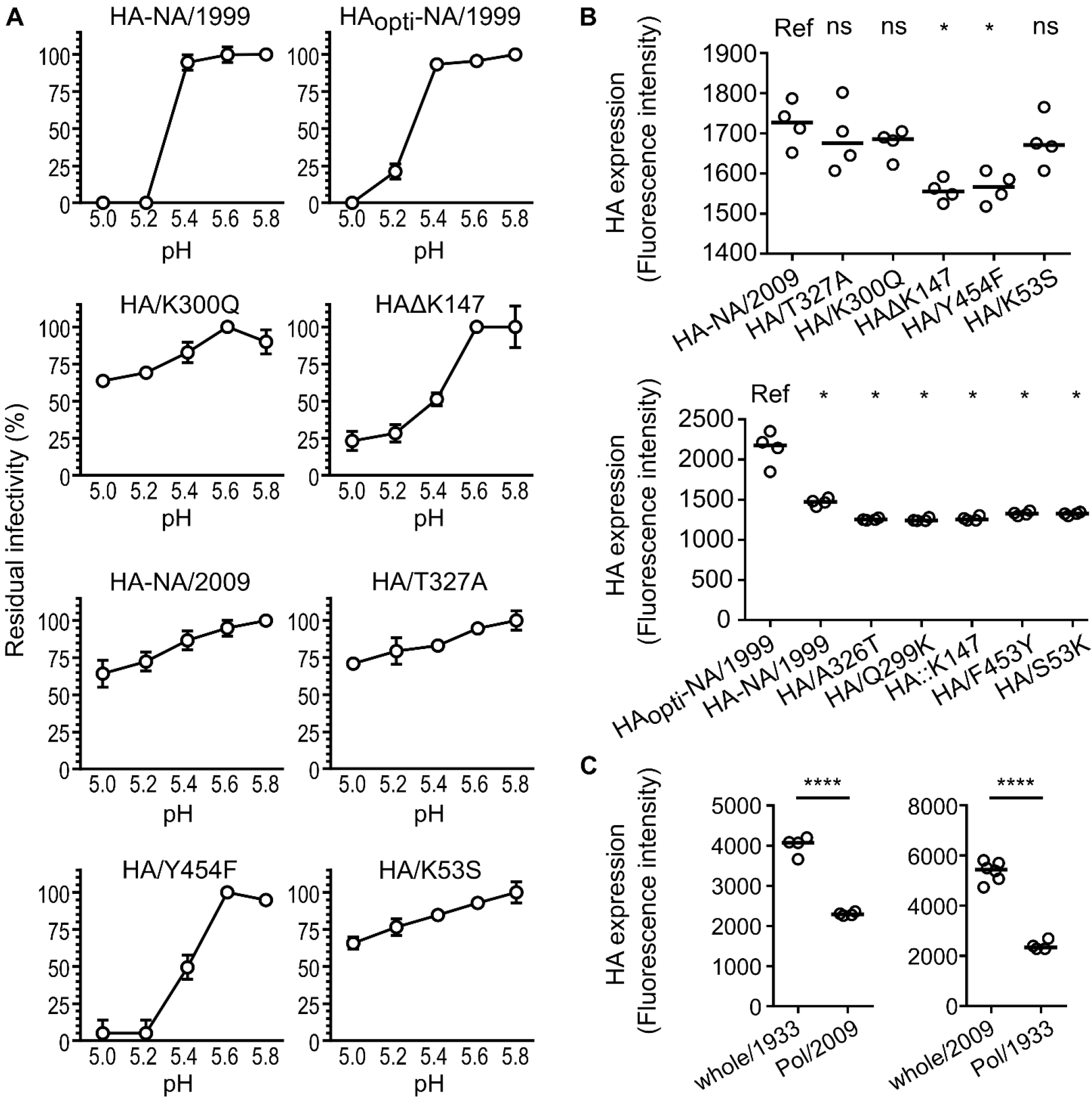
Virus stability to lower pH exposure and HA expression in infected cells. (A) H1N1 reassortant viruses either with a 1999 HA (wild-type or optimized) or a 2009 HA [wild-type or with a single amino-acid substitution) were exposed to pH-adjusted PBS ouffer for one hour and the residual infectivity values (in percentage from the initial TCID_50_ titre) were calculated. Spheres represent the mean residual infectivity and vertical lines represent the standard deviation (N=3). (B) and (C), HA surface expression in infectedMDCK cells analysed by flow cytometry detecting Alexafluor488 signal 24 hours after infection. Horizontal lines represent the median of fluorescence intensities and HA surface expression levels were compared using a Wilcoxon-Mann-Whitney test (NS, *P*>0.05; \**P*<0.02; \*\*\*\**P*<0.0001).

### Molecular determinants of the NA driving environmental persistence

Based on the results obtained with our reassortant viruses, the NA might be as important as the HA in driving the persistence phenotype (Fig. 1B). Previous studies have shown that the NA stability is strain dependent, and is linked to the calcium binding by the protein (Baker and Gandhi 1976; Burmeister, Cusack, and Ruigrok 1994). The 2009 pandemic NA has three calcium binding sites (Air 2012), as well as the 1918 NA (Xu et al. 2008), and thus presumably the 1999 NA. After alignment of the 1999 and 2009 NA protein sequences (Fig. 5A), we observed differences in the amino acids directly involved in the second and the third calcium binding sites (Air 2012), respectively near the active site and the side chains (D381, D387, and D379) (Fig 5A). We then decided to substitute the amino acids around the positions DGAD/341/NGAN (substitutions D341N and D344N), and/or DTDSD/382/GDTNN (substitutions D382G, S385N and D386N) (Fig 5A). We generated reassortant viruses with a mutated NA either in position 341 (HA-NA (341)/1999 virus), or in position 382 (HA-NA _(382)_/1999 virus) or in both positions (HA-NA _(341/382)_/1999 virus), all with a 1999 HA and a genomic backbone from the 1933 H1N1 virus. We determined the persistence of these mutant viruses in saline water at 35°C by measuring a mean inactivation slope of 10.2 CIT_50_.day^−1^ for the HA-NA_(341)_/1999 virus, 5.6 CIT_50_.day^−1^ for the HA-NA _(382)_/1999 virus and 4.3 CIT_50_.day^−1^ for the HA-NA _(341/382)_/1999 virus. In comparison, the HA-NA/1999 virus had a mean inactivation slope of 9.9 CIT_50_.day^−1^ (Fig 5B). We thus observed a positive effect of mutations at the third calcium binding site towards long-lasting persistence for the HA-NA _(382)_/1999 virus. A cumulative effect of mutations at both calcium binding sites was also observed, leading to a significant increase of HA-NA _(341/382)_/1999 virus survival compared with HA-NA _(382)_/1999 virus. The results demonstrated that amino-acids at the calcium binding sites in the NA are important molecular determinants of virus environmental persistence. Viruses with a high environmental persistence also had a stable NA, as shown in Fig. 5C and Fig. 5D, measuring the NA activity of viruses exposed or not to saline water at 35°C. Viruses with an important environmental persistence also presented a high NA activity (Fig. 5E). Unfortunately, the NA activity for the HA-NA _(341)_/1999 virus was below our detection threshold and could not be measured.

**FIG 5.**
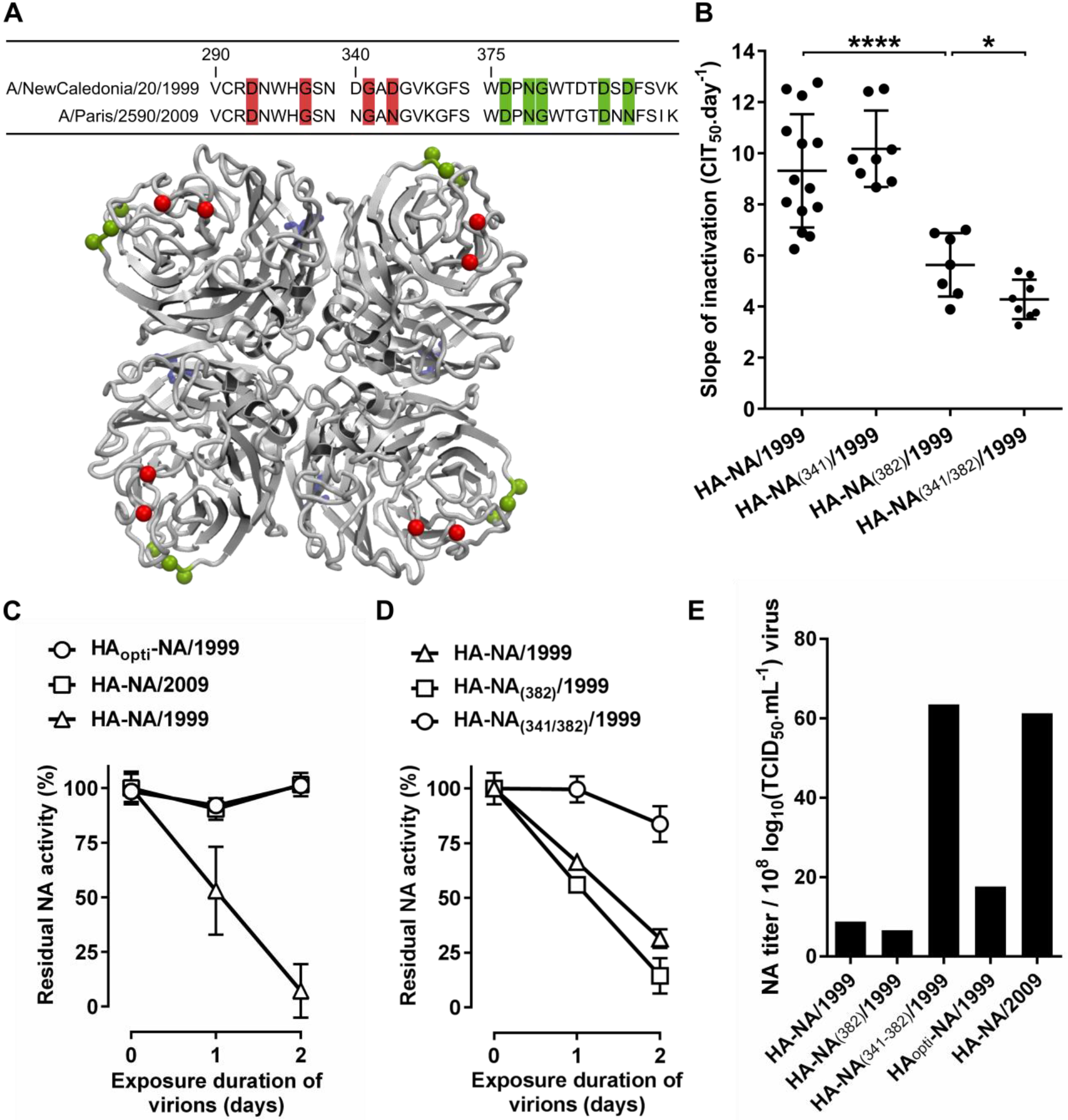
Non-synonymous substitutions in the NA change environmental persistence. (A) Alignment of amino acid sequences of NA from 1999 and 2009 H1N1 viruses. Putative amino acids involved in the second (red) and third (green) calcium-binding site are highlighted. Selected mutations on the 1999 NA are indicated by red (NGAN/341/DGAD) or green (GTDNN/382/DTDSD) spheres on the crystal structure of a 2009 H1N1 virustetrameric NA (4B7Q) (top-down view). (B) Mean inactivation slopes of viruses after exposure to saline water at 35°C. Each dot represents a single inactivation slope and their means (horizontal lines) were compared using an ANOVA test (\**P*<0.05; \*\*\*\**P*<0.0001). (C) and (D) Evolution of residual NA activity (%) during exposure of reassortant viruses in saline water at 35°C. (C) and (D) represent two individual experiments. Vertical lines represent the standard deviation (N=3). (E) NA titer of reassortant viruses measured with a two-fold serial dilution steps and then calculated for viral titer of 8 log10 (TCID_50_.mL^−1^).

## Discussion

Differences in environmental persistence among influenza virus strains of the same subtype were previously described (D.E Stallknecht and Brown 2009; Dublineau et al. 2011; Lebarbenchon et al. 2012), without providing a clear molecular basis to this phenomenon. It was also demonstrated that their viral genome was not degraded and that the viral envelope remained intact after a few days in saline water at 35°C (Dublineau et al. 2011; Shigematsu et al. 2014). The results produced in our controlled experimental model with H1N1 viral strains are the first to describe molecular basis of the IAVs persistence outside the host. Comparing the persistence of multiple engineered reassortants allowed us to identify the HA and NA proteins as major viral drivers of the virus survival outside the host (Fig. 1D and 1e and Fig. 2A).

Codon optimization of the HA gene increased considerably the viral persistence of the HA_opti_-NA/1999 virus compared with the non-optimized HA-NA/1999 virus (Fig. 2A) while they both harboured an identical HA protein. Codon usage bias that favours synonymous mutations of HA exists in nature, particularly at the more functionally constrained residues (Plotkin and Dushoff 2003). It would be interesting to investigate whether the viral codon-optimization in nature also confers adaption of the persistence phenotype to the environment. In addition, we found amino-acid changes in HA that affected the persistence of the whole virus, but without a reciprocal effect between the amino-acid changes in the 1999 or the 2009 HA. The two mutations responsible for the greatest decrease of viral persistence were located near the receptor binding site (position 147) and in the internal stem (position 454 or 453) while the others were located in the external loops of the HA1 subdomain (Fig. 3A). Some mutations affecting HA stability in low pH conditions were previously described (Russier et al. 2016; Byrd-Leotis et al. 2015) and few mutations were associated with a higher persistence outside the host (Reed et al. 2010) and to heat inactivation (Krenn et al. 2011). Similarly, we observed a link between pH stability of our 2009 HA-mutated viruses (Fig. 5A) and their persistence in water.

Our results also demonstrated that a virus with a high sensitivity to low pH still has the potential to be stable in the environment. Indeed, HA-NA/1999 and HA_opti_-NA/1999 viruses both displayed high sensitivity to low pH although they have a very different persistence phenotype in water. We also observed a higher HA surface expression levels in cells infected with HA_opti_-NA/1999 virus (Fig. 4B), in agreement with a previous study using codon optimized HA (32). Altogether, our results suggest that the survival likelihood for a virus increases when its HA surface level is higher, warranting to examine further the correlation between HA level at the cell surface and HA level at the virion surface. Results obtained with the Pol/2009 and the whole/1933 reassortant viruses, for which we expected different transcription and replication efficiencies of the HA segment, are in agreement with this finding (Fig. 4C and Fig. 1E). Although HA-NA/2009 and HA-NA/1933 are respectively more stable and less stable, compared with their parental whole/1933 and whole/2009 viruses, they have the same inactivation slope (Fig. 1D and 1E). Different replication efficiency of the polymerase complexes resulting in various HA expression levels might explain this observation.

Similarly, the less stable HAΔK147 and HA/Y454F mutant viruses (Fig. 3B) have a high sensitivity to low pH and lower HA expression level in infected cells compared to the other mutants (Fig. 4A and 4B). On the other hand, the whole/2009 and the Pol/1933 viruses had the same persistence but significant differences of HA expression levels (Fig. 4C). They both harbour the same HA of 2009, which is stable at low pH. Overall, these results suggest that the persistence of a virus with a HA resistant to low pH does not vary with HA expression level.

After losing their infectivity, HA-NA/2009 and HA-NA/1999 viruses were still able to attach to cells but could not trigger viral fusion after endocytic uptake. Three or four neighbouring HA trimers are probably mandatory to reach the hemifusion step occurring during the fusion process (33). This number might not be reached for viruses that do not have a sufficient density of functional HA at their surface and could explain the observed absence of fusion after exposure to a hostile environment. This observation might in turn provide an explanation for how viruses with higher HA surface levels stay stable longer, as the HA inactivation is probably progressive in the environment as suggested by our results on the loss hemagglutination activity (Fig. 3A).

We provided evidences that the NA protein is also a driver of IAVs environmental persistence. Our results highlighted the importance of amino acid positions in the calcium binding sites for both the NA stability and virus survival. It was previously shown that the NA from the 2009 and 1918 H1N1 pandemic viruses possess a third calcium-binding site coordinated by side chains (Xu et al. 2008; Air 2012), suggesting that all the human N1 may contain three calcium-binding sites. Based on a sequence alignment between the NA of the 2009 and 1999 H1N1 viruses and observations of the structural features, we substituted key amino acids involved in the second and the third calcium-binding sites. These mutations induced an important increase of the environmental persistence of both the HA-NA _(382)_/1999 and HA-NA _(341/382)_/1999 viruses compared with the HA-NA/1999 virus (Fig. 5B). Our results raised new questions on this putative third calcium-binding site in the human N1, which may not be present or possess a weaker affinity for calcium ions in the NA of the pre-pandemic H1N1 of 1999. Moreover, we observed that the more stable virus in the environment, such as HA_opti_-NA/1999 virus and the double mutant HA-NA _(341/382)_/1999 virus, induced a high HA surface expression level or had a stable and high NA activity (Fig. 4B and Fig. 5E). The mutations introduced in the HA or the NA probably modified the viral HA/NA balance, which might be an important molecular determinant of environmental persistence, possibly by increasing sialic acid binding, which was suggested to stabilize the influenza virus (36, 37). These results raised new questions about the role of the NA protein in the viral entry, which is not yet fully understood.

Based on previous published results, we found that a log-linear model drives influenza A virus inactivation in our experimental model (Dublineau et al. 2011). We observed that the distributions of the inactivation slopes among the different viruses were significantly different (Bartlett’s statistic, P<0.05), so we used a non-parametric Wilcoxon-Mann-Whitney test to compare virus persistence. Moreover, we observed that the standard deviation of the mean inactivation slopes tends to increase in the experiments performed with low persistence viruses compared with experiments with high persistence viruses (Fig. S3). It is possibly because of minor genetic variants or morphological variants, present in every viral population, which could have a higher persistence than the average population. Minor variants proportion may increase during the viral decay monitoring of low persistence viruses. Indeed, the average population of high persistence viruses remained infectious for a longer period and is more homogeneous after a long time at 35°C compared with low persistence viruses that are submitted to a more important bottleneck.

The putative role of particle persistence in the environment for IAV transmission has been discussed in reviews (39, 40). Using computational approaches, it was suggested that the virus persistence in the environmental reservoir is an important parameter impacting overwintering of IAVs, infection probability of migratory ducks in low density population areas as well as spatial variations of IAV spreading (9, 41, 42). IAVs persistence could also explain their evolutionary dynamic. It was shown that hemagglutinin diversity found in avian IAVs is positively correlated to viral persistence (Roche et al. 2014). This could allow viral strains to adapt to local environmental conditions (Lebarbenchon et al. 2012). In addition, evidence of IAV persistence was shown in Antarctica (43) and in Siberian ice lakes (44) where viruses are most likely stable for years (Dublineau et al. 2011; Brown et al. 2007), justifying to further investigate the role of long-term persistence on virus reintroduction. The present study provides for the first time experimental data highlighting the role of the HA variability in driving influenza persistence in the environment.

In conclusion, the molecular drivers of influenza virus persistence that we identified in the present study could help refining ecological model of IAVs transmission and their genetic diversity in the environment. Our results are providing an experimental basis to further investigate the role of the HA and NA proteins in driving the phenotype of persistence among all influenza viral subtypes and establish the impact of their persistence on viral transmission in the environment.

## Methods

### Cells and viruses

Madin-Darby canine kidney epithelial (MDCK) cells were maintained in Modified Eagle’s Medium (MEM) (GIBCO, ThermoFisher Scientific), supplemented with 10% foetal calf serum (FCS) and antibiotics (100 units.mL^−1^ penicillin, 100 mg.mL^−1^ streptomycin, GIBCO, Life Technologies). Human embryonic kidney cells 293T (HEK-293T) were maintained in Dulbecco’s Modified Eagle’s medium (DMEM) (GIBCO, ThermoFisher Scientific) supplemented with 10% FCS. All cells were incubated at 37°C in humidified 5% CO2 incubator. MDCK cells were infected at a multiplicity of infection (MOI) of 10^−4^ plaque-forming units per cell (pfu.cell^−1^) and maintained in MEM without FCS in the presence of 1 μg.mL^−1^ of TPCK-trypsin (Trypsin/L-1-Tosylamide-2-phenylethyl chloromethyl Ketone, Whortington Biochemical Corporation) at 35°C for 3 days to generate stock viruses. The clarified supernatants were harvested, aliquoted and stored at-80°C.

### Generation of reassortant viruses and mutated viruses

Recombinant viruses were rescued after co-transfection of MDCK and 293T co-cultivated cells using FuGENE HD transfection reagent (Promega) at a ratio of 3:1 (μL:μg). A/WSN/1933 recombinant virus was rescued by reverse genetics using a twelve plasmids transfection system, with eight pPol transcription plasmids containing an individual genomic segment under the control of a truncated human RNA polymerase I promoter and upstream the hepatitis Delta virus ribozyme as well as four pCDNA expression plasmids containing either the NP, PA, PB1 or PB2 gene under the control of a cytomegalovirus promoter as described previously (45). A/Bretagne/7608/2009 recombinant virus was rescued by reverse genetics using an 8 plasmids transfection system as described previously (Hoffmann et al. 2000). Reassortant viruses between A/WSN/1933 and A/Bretagne/7608/2009 strains were rescued using plasmids from those two reverse genetics systems (Reassortant viruses are listed in table 1). Reassortant viruses harbouring HA and NA from A/Paris/2590/2009 or A/NewCaledonia/20/1999 strains with the internal genes of A/WSN/1933 were rescued using the twelve plasmids transfection system for A/WSN/1933. Nucleotide changes were introduced into the pPol-HA plasmid by using the QuikChange II site-directed mutagenesis kit (Agilent) in accordance with the manufacturer’s instructions. Because the pPol-HA plasmid expressing the A/NewCaledonia/20/1999 HA segment was unstable in bacteria, the HA DNA sequence was replaced by a codon-optimized sequence and used for transfection. Virus identity and absence of unintended mutations were confirmed by Sanger sequencing using BigDye Terminator v1.1 cycle sequencing kit (Applied Biosystems) and a 3730 DNA Analyzer (Applied Biosystems).

### Virus persistence in saline water at 35°C

Virus persistence in saline water at 35°C was studied as follow: viral suspensions were diluted 10 times in saline distilled water (35 g.L^−1^ NaCl) and placed in a humidified incubator (5% CO2, 35°C) for 1 hour, 24 hours or 48 hours. The pH of the saline water did not vary between experiments. In order to quantify their residual infectivity, MDCK cells were seeded on a 16-well microtiter plate (30000 cells per well) coated with microelectrode sensors in the xCELLigence^®^ Real-Time Cell Analysis (RTCA) DP instrument (ACEA Bioscience, Inc.) and grown for 24 hours (5% CO_2_, 37°C). Cells were then infected by exposed viruses and cell impedance, expressed as an arbitrary unit called the Cell Index (CI), was measured through the electrodes every 15 min. Cell Index decrease due to virus-induced cytopathogenic effect was quantified with the CIT_50_ value, which is the necessary time in hours to measure a 50% decrease from the initial CI, always set as the CI value 5 hours after infection. CIT_50_ values are linearly correlated to the TCID_50_ titres of the viral suspensions (47) (Fig. S4). Thus, the increase of the CIT_50_ values over time reflects the loss of infectivity. For each exposed viral suspension, CIT_50_ values were obtained at different exposure times and plotted to calculate a linear regression slope referred to as the inactivation slope. All data shown on viral persistence have been obtained at different periods and with different viral stock productions. Each replicate value of the inactivation slopes refers to a replicate virus suspension tested for persistence.

### Virus concentration for functional analyses

For the experiments described below, harvested supernatants of stock viruses were clarified and concentrated using a vivaspin 20 centrifugal concentrator (1 000 000 MWCO, Sartorius), to reduce the initial volume by 10 times. Concentrated supernatants were then diluted in saline distilled water (35g.L^−1^ NaCl) at a ratio of 1:10 and placed either at 4°C (0 day exposure) or at 35°C for 5 days and then kept at 4°C until used in further analysis.

### Hemagglutination assay

Exposed viruses were diluted in two-fold dilution steps with PBS in a 96-wells plate and mixed with an equal volume of a 0.75% suspension of fresh guinea pig erythrocytes. The mixture was incubated for 1 hour at room temperature before observing erythrocytes aggregation.

### Neuraminidase activity assay

Neuraminidase activity of exposed viruses was measured using a NA-Fluor kit (Applied Biosystems) according to manufacturer’s protocol. Viruses were diluted in two-fold dilution steps and mixed with an equal volume of MUNANA (2′-(4-methylumbelliferyl)-alpha-D-N-acetylneuraminic acid) substrate. After 1 or 2 hours, the level of fluorescence was quantified with a microplate fluorimeter (TriStar^2^ LB942, Berthold).

### Fluorescence microscopy

For influenza virus nucleoprotein (NP) labelling, MDCK cells were seeded on glass coverslips in 6-well plates for 24 hours before inoculation with exposed viruses diluted 1:10 in MEM. Twenty minutes or 2 hours post infection, cells were washed twice with MEM and fixed with paraformaldehyde 2% (w/v) for 10 min. Cell membranes were labelled using Alexa Fluor 488 conjugated wheat germ agglutinin (ThermoFisher Scientific) at 4 μg.mL^−1^ for 10 min and permeabilized during 20 min in PBS buffer containing 0.2% Triton X-100. Fixed cells were incubated at 4°C overnight in the presence of an anti-NP antibody (MAB8257, Merck Millipore) diluted in PBS buffer containing 2% BSA at a concentration of 1μg.mL^−1^, then incubated at room temperature for 2 hours with an Alexa Fluor 647 F (ab′)2-Goat anti-Mouse IgG (H+L) Cross-Adsorbed Secondary antibody (ThermoFisher Scientific) diluted at a concentration of 2 μg.mL^−1^. For Influenza virus envelope labelling and endosomal fusion analyses, concentrated viral suspensions were incubated in the presence of rhodamine B (R18, ThermoFisher Scientific) and Dioc_18_ (3) (3,3′-Dioctadecyloxacarbocyanine Perchlorate, ThermoFisher Scientific) for 2 hours at room temperature both at a final concentration of 20 μM. Viral suspensions were then applied on a PD-10 desalting column (GE Healthcare) using gravity flow according to the manufacturer’s instructions to remove fluorophore in suspension. MDCK cells seeded on glass coverslips in 6-well plates for 24 hours were placed at 4°C for 5 min prior to infection in order to synchronize viral fusion (Schelker et al. 2016) and were then inoculated with labelled viral suspensions. After infection, the cells were incubated at 35°C for 20 min and then washed twice with MEM and fixed with paraformaldehyde 2% (w/v) for 10 min. In all experiments, cell nuclei were labelled using Hoescht 33342 diluted in PBS at a concentration of 5 μg.mL^−1^ and incubated for 10 min. Glass coverslips were then mounted on a glass slide with Prolong gold antifade reagent (ThermoFisher Scientific). Confocal laser scanning of fluorescence was performed using LSM700 inverted microscope (Zeiss) equipped with a plan apochromat 10X objective with a numerical aperture (N.A) of 0.45 and an enhanced contrast plan-neofluar 40X oil objective with a N.A of 1.3.

### HA stability to pH

Three independent virus replicates were diluted in a pH-controlled PBS solution (at a resolution of 0.2) with citric acid and incubated 1 hour at room temperature. Residual infectivity of each replicate was then quantified by a TCID50 method, as described previously (Dublineau et al. 2011).

### HA expression level by flow-cytometry

Cell surface HA expression of infected cells was measured by immunofluorescence using flow cytometry. MDCK cells were infected for 24 hours in 6-well plates at a MOI of 0.5 pfu.cell^−1^ in the presence of TPCK-trypsin. Cell monolayers were washed twice with PBS, re-suspended using trypsin-EDTA (ThermoFisher Scientific), fixed with paraformaldehyde 2% (w/v) for 10 min and incubated with an anti-HA monoclonal antibody (Sinobiological) at a concentration of 1 μg.mL^−1^ at 4°C overnight. Cells were then incubated for 30 min at room temperature in the presence of a species-specific Alexafluor 488 conjugated secondary antibody (ThermoFisher Scientific). After centrifugation, cells were re-suspended in PBS and analysed with an Attune NxT Flow Cytometer (ThermoFisher Scientific). For each replicate, 50,000 cells were analysed with the same parameters.

### Statistical analyses and software

Numerical data were analysed with Prism software version 6.07 (Graph Pad Software). If needed, a Grubb’s test was performed to identify and remove outliers. ANOVA tests were performed for statistical comparison when variances were equal across samples (Bartlett’s test). Otherwise a two-tailed, Wilcoxon-Mann-Whitney’s test, was used to compare samples distribution. DNA sequence alignments and Sanger sequencing analysis were performed using CLC Main Workbench version 7.7.3 (Qiagen). Hemagglutinin structure was visualized using the PyMOL Molecular Graphics system version 1.8 (Schrodinger, LLC). Microscopy imaging data were processed using LSM software Zen Blue edition version 2.3 (Zeiss) and flow cytometry data were analysed with FlowJo (LLC) software.

### Data availability

The data that support the findings of this study are available from the corresponding author upon request.

## Acknowledgements

We thank Olivier Sawoo (Environment and Infectious Risk Unit) for the pre-screen of HA mutants based on lentiviral pseudo-particles. We thank Cyril Barbezange and Sandie Munier (laboratory Molecular Genetics of RNA Viruses, Institut Pasteur Paris) for providing reagents and advices. We are also grateful to Gaëlle Lelandais (Institut de Biologie Intégrative de la Cellule, CNRS UMR 9198) for statistical assistance and to Nathalie Pardigon (Environment and Infectious Risk Unit) for assistance with the manuscript.

This work was supported by the European funded PREDEMICS program (Preparedness, Prediction, and Prevention of Emerging Zoonotic Viruses with Pandemic Potential using Multidisciplinary Approaches) FP7/2007-2013 under Grant Agreement No 278433, the IBEID labex as well as BNP Paribas.

## References

Air, Gillian M. 2012. “Influenza Neuraminidase.” Influenza and Other Respiratory Viruses 6 (4): 245–56. https://doi.org/10.1111/j.1750-2659.2011.00304.x.

Bajimaya, Shringkhala, Tünde Frankl, Tsuyoshi Hayashi, and Toru Takimoto. 2017. “Cholesterol Is Required for Stability and Infectivity of Influenza A and Respiratory Syncytial Viruses.” Virology 510 (July): 234–41. https://doi.org/10.1016/j.virol.2017.07.024.

Baker, N. J., and S. S. Gandhi. 1976. “Effect of Ca++ on the Stability of Influenza Virus Neuraminidase.” Archives of Virology 52 (1-2): 7–18.

Brown, Justin D., Ginger Goekjian, Rebecca Poulson, Steve Valeika, and David E. Stallknecht. 2009. “Avian Influenza Virus in Water: Infectivity Is Dependent on PH, Salinity and Temperature.” Veterinary Microbiology 136 (1): 20–26. https://doi.org/10.1016/j.vetmic.2008.10.027.

Brown, Justin D., David E. Swayne, Robert J. Cooper, Rachel E. Burns, and David E. Stallknecht. 2007. “Persistence of H5 and H7 Avian Influenza Viruses in Water.” Avian Diseases 51 (s1): 285–89. https://doi.org/10.1637/7636-042806R.1.

Burmeister, Wilhelm Pascal, Stephen Cusack, and Rob W. H. Ruigrok. 1994. “Calcium Is Needed for the Thermostability of Influenza B Virus Neuraminidase.” Journal of General Virology 75 (2): 381–88. https://doi.org/10.1099/0022-1317-75-2-381.

Byrd-Leotis, Lauren, Summer E. Galloway, Evangeline Agbogu, and David A. Steinhauer. 2015. “Influenza Hemagglutinin (HA) Stem Region Mutations That Stabilize or Destabilize the Structure of Multiple HA Subtypes.” Journal of Virology 89 (8): 4504–16. https://doi.org/10.1128/JVI.00057-15.

Chu, C. M. 1948. “Inactivation of Haemagglutinin and Infectivity of Influenza and Newcastle Disease Viruses by Heat and by Formalin.” The Journal of Hygiene 46 (3): 247–51.

Dublineau, Amélie, Christophe Batéjat, Anthony Pinon, Ana Maria Burguière, India Leclercq, and Jean-Claude Manuguerra. 2011. “Persistence of the 2009 Pandemic Influenza A (H1N1) Virus in Water and on Non-Porous Surface.” PLoS ONE 6 (11): e28043. https://doi.org/10.1371/journal.pone.0028043.

Fang, Ying, Peifang Ye, Xiaobo Wang, Xiao Xu, and William Reisen. 2011. “Real-Time Monitoring of Flavivirus Induced Cytopathogenesis Using Cell Electric Impedance Technology.” Journal of Virological Methods 173 (2): 251–58. https://doi.org/10.1016/j.jviromet.2011.02.013.

Fodor, Ervin, Louise Devenish, Othmar G. Engelhardt, Peter Palese, George G. Brownlee, and Adolfo García-Sastre. 1999. “Rescue of Influenza A Virus from Recombinant DNA.” Journal of Virology 73 (11): 9679–82.

Hirose, Ryohei, Takaaki Nakaya, Yuji Naito, Tomo Daidoji, Yohei Watanabe, Hiroaki Yasuda, Hideyuki Konishi, and Yoshito Itoh. 2017. “Mechanism of Human Influenza Virus RNA Persistence and Virion Survival in Feces: Mucus Protects Virions From Acid and Digestive Juices.” The Journal of Infectious Diseases 216 (1): 105–9. https://doi.org/10.1093/infdis/jix224.

Hoffmann, Erich, Gabriele Neumann, Yoshihiro Kawaoka, Gerd Hobom, and Robert G. Webster. 2000. “A DNA Transfection System for Generation of Influenza A Virus from Eight Plasmids.” Proceedings of the National Academy of Sciences 97 (11): 6108–13. https://doi.org/10.1073/pnas.100133697.

Hurt, Aeron C., Yvonne C. F. Su, Malet Aban, Heidi Peck, Hilda Lau, Chantal Baas, Yi-Mo Deng, et al. 2016. “Evidence for the Introduction, Reassortment, and Persistence of Diverse Influenza A Viruses in Antarctica.” Journal of Virology 90 (21): 9674–82. https://doi.org/10.1128/JVI.01404-16.

Imai, Masaki, Tokiko Watanabe, Masato Hatta, Subash C. Das, Makoto Ozawa, Kyoko Shinya, Gongxun Zhong, et al. 2012. “Experimental Adaptation of an Influenza H5 HA Confers Respiratory Droplet Transmission to a Reassortant H5 HA/H1N1 Virus in Ferrets.” Nature 486 (7403): 420–28. https://doi.org/10.1038/nature10831.

Ivanovic, Tijana, Jason L Choi, Sean P Whelan, Antoine M van Oijen, and Stephen C Harrison. 2013. “Influenza-Virus Membrane Fusion by Cooperative Fold-Back of Stochastically Induced Hemagglutinin Intermediates.” ELife 2 (February). https://doi.org/10.7554/eLife.00333.

Jiang, Yongping, Kangzhen Yu, Hongbo Zhang, Pingjing Zhang, Chenjun Li, Guobin Tian, Yanbing Li, et al. 2007. “Enhanced Protective Efficacy of H5 Subtype Avian Influenza DNA Vaccine with Codon Optimized HA Gene in a PCAGGS Plasmid Vector.” Antiviral Research 75 (3): 234–41. https://doi.org/10.1016/j.antiviral.2007.03.009.

Keeler, Shamus P., Melinda S. Dalton, Alan M. Cressler, Roy D. Berghaus, and David E. Stallknecht. 2014. “Abiotic Factors Affecting the Persistence of Avian Influenza Virus in Surface Waters of Waterfowl Habitats.” Applied and Environmental Microbiology 80 (9): 2910–17. https://doi.org/10.1128/AEM.03790-13.

Krenn, Brigitte M., Andrej Egorov, Ekaterina Romanovskaya-Romanko, Markus Wolschek, Sabine Nakowitsch, Tanja Ruthsatz, Bettina Kiefmann, et al. 2011. “Single HA2 Mutation Increases the Infectivity and Immunogenicity of a Live Attenuated H5N1 Intranasal Influenza Vaccine Candidate Lacking NS1.” PLOS ONE 6 (4): e18577. https://doi.org/10.1371/journal.pone.0018577.

Lang, Andrew S., Anke Kelly, and Jonathan A. Runstadler. 2008. “Prevalence and Diversity of Avian Influenza Viruses in Environmental Reservoirs.” Journal of General Virology 89 (2): 509–19. https://doi.org/10.1099/vir.0.83369-0.

Lebarbenchon, Camille, Chris J. Feare, François Renaud, Frédéric Thomas, and Michel Gauthier-Clerc. 2010. “Persistence of Highly Pathogenic Avian Influenza Viruses in Natural Ecosystems.” Emerging Infectious Diseases 16 (7): 1057–62. https://doi.org/10.3201/eid1607.090389.

Lebarbenchon, Camille, Srinand Sreevatsan, Thierry Lefèvre, My Yang, Muthannan A. Ramakrishnan, Justin D. Brown, and David E. Stallknecht. 2012. “Reassortant Influenza A Viruses in Wild Duck Populations: Effects on Viral Shedding and Persistence in Water.” Proceedings of the Royal Society B: Biological Sciences 279 (1744): 3967–75. https://doi.org/10.1098/rspb.2012.1271.

Millero, Frank J., Rainer Feistel, Daniel G. Wright, and Trevor J. McDougall. 2008. “The Composition of Standard Seawater and the Definition of the Reference-Composition Salinity Scale.” Deep Sea Research Part I: Oceanographic Research Papers 55 (1): 50–72. https://doi.org/10.1016/j.dsr.2007.10.001.

Nazir, Jawad, Renate Haumacher, Anthony Ike, Petra Stumpf, Reinhard Böhm, and Rachel E. Marschang. 2010. “Long-Term Study on Tenacity of Avian Influenza Viruses in Water (Distilled Water, Normal Saline, and Surface Water) at Different Temperatures.” Avian Diseases 54 (s1): 720–24. https://doi.org/10.1637/8754-033109-ResNote.1.

Plotkin, Joshua B., and Jonathan Dushoff. 2003. “Codon Bias and Frequency-Dependent Selection on the Hemagglutinin Epitopes of Influenza A Virus.” Proceedings of the National Academy of Sciences 100 (12): 7152–57. https://doi.org/10.1073/pnas.1132114100.

Pohl, Marie O., Thomas O. Edinger, and Silke Stertz. 2014. “Prolidase Is Required for Early Trafficking Events during Influenza A Virus Entry.” Journal of Virology 88 (19): 11271–83. https://doi.org/10.1128/JVI.00800-14.

Poulson, R. L., S. M. Tompkins, R. D. Berghaus, J. D. Brown, and D. E. Stallknecht. 2016. “Environmental Stability of Swine and Human Pandemic Influenza Viruses in Water under Variable Conditions of Temperature, Salinity, and PH.” Applied and Environmental Microbiology 82 (13): 3721–26. https://doi.org/10.1128/AEM.00133-16.

Reed, Mark L., Olga A. Bridges, Patrick Seiler, Jeong-Ki Kim, Hui-Ling Yen, Rachelle Salomon, Elena A. Govorkova, Robert G. Webster, and Charles J. Russell. 2010. “The PH of Activation of the Hemagglutinin Protein Regulates H5N1 Influenza Virus Pathogenicity and Transmissibility in Ducks.” Journal of Virology 84 (3): 1527–35. https://doi.org/10.1128/JVI.02069-09.

Roche, Benjamin, John M. Drake, Justin Brown, David E. Stallknecht, Trevor Bedford, and Pejman Rohani. 2014. “Adaptive Evolution and Environmental Durability Jointly Structure Phylodynamic Patterns in Avian Influenza Viruses.” PLOS Biology 12 (8): e1001931. https://doi.org/10.1371/journal.pbio.1001931.

Russier, Marion, Guohua Yang, Jerold E. Rehg, Sook-San Wong, Heba H. Mostafa, Thomas P. Fabrizio, Subrata Barman, et al. 2016. “Molecular Requirements for a Pandemic Influenza Virus: An Acid-Stable Hemagglutinin Protein.” Proceedings of the National Academy of Sciences of the United States of America 113 (6): 1636–41. https://doi.org/10.1073/pnas.1524384113.

Sawoo, Olivier, Amélie Dublineau, Christophe Batéjat, Paul Zhou, Jean-Claude Manuguerra, and India Leclercq. 2014. “Cleavage of Hemagglutinin-Bearing Lentiviral Pseudotypes and Their Use in the Study of Influenza Virus Persistence.” PLoS ONE 9 (8): e106192. https://doi.org/10.1371/journal.pone.0106192.

Schelker, Max, Caroline Maria Mair, Fabian Jolmes, Robert-William Welke, Edda Klipp, Andreas Herrmann, Max Flöttmann, and Christian Sieben. 2016. “Viral RNA Degradation and Diffusion Act as a Bottleneck for the Influenza A Virus Infection Efficiency.” PLOS Comput Biol 12 (10): e1005075. https://doi.org/10.1371/journal.pcbi.1005075.

Shigematsu, Sayuri, Amélie Dublineau, Olivier Sawoo, Christophe Batéjat, Toshifumi Matsuyama, India Leclercq, and Jean-Claude Manuguerra. 2014. “Influenza A Virus Survival in Water Is Influenced by the Origin Species of the Host Cell.” Influenza and Other Respiratory Viruses 8 (1): 123–30. https://doi.org/10.1111/irv.12179.

Sooryanarain, Harini, and Subbiah Elankumaran. 2015. “Environmental Role in Influenza Virus Outbreaks.” Annual Review of Animal Biosciences 3 (1): null. https://doi.org/10.1146/annurev-animal-022114-111017.

Souris, Marc, Daniel Gonzalez, Witthawat Wiriyarat, Kamlang Chumpolbanchorn, Supaluk Khaklang, Suwannapa Ninphanomchai, Weena Paungpin, et al. 2015. “Potential Role of Fresh Water Apple Snails on H5N1 Influenza Virus Persistence and Concentration in Nature.” Air & Water Borne Diseases 4 (1). https://doi.org/10.4172/2167-7719.1000119.

Stallknecht, David E., Virginia H. Goekjian, Benjamin R. Wilcox, Rebecca L. Poulson, and Justin D. Brown. 2010. “Avian Influenza Virus in Aquatic Habitats: What Do We Need to Learn?” Avian Diseases 54 (1 Suppl): 461–65. https://doi.org/10.1637/8760-033109-Reg.1.

Stallknecht, D.E, and J. D. Brown. 2009. “Tenacity of Avian Influenza Viruses.” Revue Scientifique et Technique de I’OIE 28 (1): 59–67.

Vittecoq, Marion, Hermann Gauduin, Thibault Oudart, Olivier Bertrand, Benjamin Roche, Matthieu Guillemain, and Olivier Boutron. 2017. “Modeling the Spread of Avian Influenza Viruses in Aquatic Reservoirs: A Novel Hydrodynamic Approach Applied to the Rhône Delta (Southern France).” Science of The Total Environment 595 (October): 787–800. https://doi.org/10.1016/j.scitotenv.2017.03.165.

Weber, Thomas P, and Nikolaos I Stilianakis. 2008. “Inactivation of Influenza A Viruses in the Environment and Modes of Transmission: A Critical Review.” The Journal of Infection 57 (5): 361–73. https://doi.org/10.1016/j.jinf.2008.08.013.

Xu, Xiaojin, Xueyong Zhu, Raymond A. Dwek, James Stevens, and Ian A. Wilson. 2008. “Structural Characterization of the 1918 Influenza Virus H1N1 Neuraminidase.” Journal of Virology 82 (21): 10493–501. https://doi.org/10.1128/JVI.00959-08.

Zhang, Gang, Dany Shoham, David Gilichinsky, Sergei Davydov, John D. Castello, and Scott O. Rogers. 2006. “Evidence of Influenza A Virus RNA in Siberian Lake Ice.” Journal of Virology 80 (24): 12229–35. https://doi.org/10.1128/JVI.00986-06.

Zhang, Hongbo, Yan Li, Jianjun Chen, Quanjiao Chen, and Ze Chen. 2014. “Perpetuation of H5N1 and H9N2 Avian Influenza Viruses in Natural Water Bodies.” Journal of General Virology 95 (7): 1430–35. https://doi.org/10.1099/vir0.063438-0.

